# Rapid estimates of leaf litter chemistry using reflectance spectroscopy

**DOI:** 10.1101/2023.11.27.568939

**Authors:** Shan Kothari, Sarah E. Hobbie, Jeannine Cavender-Bares

## Abstract

Measuring the chemical traits of leaf litter is important for understanding plants’ roles in nutrient cycles, including through nutrient resorption and litter decomposition, but conventional leaf trait measurements are often destructive and labor-intensive. Here, we develop and evaluate the performance of partial least-squares regression (PLSR) models that use reflectance spectra of intact or ground leaves to estimate leaf litter traits, including carbon and nitrogen concentration, carbon fractions, and leaf mass per area (LMA). Our analyses included more than 300 samples of senesced foliage from 11 species of temperate trees, including needleleaf and broadleaf species. Across all samples, we could predict each trait with moderate-to-high accuracy from both intact-leaf litter spectra (validation *R^2^* = 0.543-0.941; %RMSE = 7.49-18.5) and ground-leaf litter spectra (validation *R^2^* = 0.491-0.946; %RMSE = 7.00-19.5). Notably intact-leaf spectra yielded better predictions of LMA. Our results support the feasibility of building models to estimate multiple chemical traits from leaf litter of a range of species. In particular, the success of intact-leaf spectral models allows non-destructive trait estimation in a matter of seconds, which could enable researchers to measure the same leaves over time in studies of nutrient resorption.

## Introduction

Long-lived plants resorb and store a large fraction of the nutrients from their leaves before they senesce. These reserves support early growth in the following growing season and reduce the need for nutrient uptake from the soil (El Zein et al. 2011). But nutrient resorption (much like soil nutrient uptake) has metabolic and physiological costs, so the optimal level of nutrient resorption varies among environments (Wright & Westoby 2003). Accordingly, species vary broadly in their nutrient resorption efficiency—the fraction of a given nutrient that is resorbed (Vergutz et al. 2012). For example, plants often (Kobe et al. 2005; Hayes et al. 2014; Yuan & Chen 2015), though not always (Diehl et al. 2003), resorb a greater fraction of leaf nutrients when soil nutrients are scarce. This adjustment of resorption efficiency in response to the environment highlights the key role of resorption in plant nutrient economics.

Resorption often leaves the resulting leaf litter relatively nutrient-poor, which can limit the rate of decomposition of newly shed litter by soil microorganisms (Berg 2014). Hence, nutrient-poor litter decomposes more slowly than nutrient-rich litter at first (Cornwell et al. 2008), although this trend can reverse in later stages (Prescott 2010; Berg 2014; Gill et al. 2021). Because litter nutrient levels influence the nutrient flux into soil inorganic or microbial pools via decomposition, resorption may be a key process for explaining how plants influence their environment (Hobbie 2015). Despite the metabolic changes that occur during senescence, relationships between fresh-leaf and leaf litter composition may help explain why soil processes can often be inferred from remote sensing of aboveground vegetation (Madritch et al. 2020, Cavender-Bares et al. 2022).

Addressing many ecological questions about nutrient resorption and litter decomposition requires measuring litter chemistry. Because measuring resorption efficiency requires the nutrient content of leaves both before and after senescence, it can be expensive and labor-intensive. Furthermore, because measuring nutrient concentration through elemental analysis is destructive, the same exact leaves cannot be measured at each step. Decomposition also depends on litter traits beyond macronutrient concentrations, including lignin, cellulose, condensed tannins, and various micronutrients (Schweitzer et al. 2004; Cornwell et al. 2008; Talbot & Treseder 2012; Keiluweit et al. 2015; Bourget et al. 2023). The number of traits and samples can add up to make even simple projects expensive and time-consuming.

Such complications may make it harder to understand and predict the fluxes of carbon and mineral nutrients between plants, litter, and soil—the determinants of which are still far from well-understood. For example, large-scale comparative tests of the role of nutrient resorption in plant life history or nutrient economics are still rare (e.g., Freschet et al. 2010; Rea et al. 2018). Meanwhile, research continues to reveal new paths through which a plant community’s litter traits or diversity interact with microbial communities and the abiotic environment to influence decomposition rates (Liu et al. 2020; Grossman et al. 2020; Gill et al. 2021; Bourget et al. 2023). Fast, simple estimates of litter traits could make it easier both to study the functional significance of nutrient resorption and to estimate litter decomposability rapidly across taxa and ecosystems.

The need for faster trait estimates has led some plant ecologists to turn to reflectance spectroscopy, a technique that involves measuring the reflectance of light from a sample across many wavelengths. Many kinds of samples, including leaf tissue, are analyzed using the 400-2400 nm range. Reflectance in this range can be used to estimate many leaf structural and chemical traits because those traits influence how leaves scatter and absorb light across wavelengths.

Many widely studied functional traits nevertheless cannot be identified with specific structures or chemical constituents that have unique absorption features. Leaf nitrogen concentration (N_mass_), for example, includes nitrogen in molecules as varied as RuBisCO and chlorophyll. This challenge is among the reasons that many plant scientists have adopted a multivariate empirical approach to linking traits and reflectance spectra using statistical techniques like partial least-squares regression (PLSR; Burnett et al. 2021). Researchers have built PLSR models predicting traits like N_mass_ and leaf mass per area (LMA) with a high degree of accuracy from spectra, measured either while the leaf was fresh or after drying and grinding it into powder (Serbin et al. 2014; Serbin et al. 2019; Kothari et al. 2023a; Kothari et al. 2023b). Further research has shown that spectral models can predict traits as varied as defense compounds (Couture et al. 2016), water status (Cotrozzi et al. 2017; Kothari et al. 2023a), leaf age (Chavana-Bryant et al. 2017), and photosynthetic capacity (Dechant et al. 2017).

The vast majority of this research is based on measurements taken during the growing season, before the onset of leaf senescence. These models are not expected to accurately predict the traits of senesced leaves, which are well outside the scope of their training data. Some researchers have built models to predict decomposition-related traits (Joffre et al. 1992; McTiernan et al. 2003; Coûteaux et al. 2005; Hobbie 2005; Parsons et al. 2011; Petit Bon et al. 2020) or the decomposition rate itself (Gillon et al. 1999; Fortunel et al. 2009; Parsons et al. 2011) from ground leaf litter using near-infrared (NIR) reflectance spectroscopy. Many traits that influence decomposition also influence forage quality, and there is an extensive body of research predicting forage traits using NIR spectroscopy, including in senescent foliage (e.g., Norris et al. 1976). Such studies have often shown high accuracy (*R^2^* > 0.9) in predicting traits like N_mass_, carbon fractions, and phenolics, albeit usually using a limited number (<120) of calibration samples from one or a few species.

Here, we expand on prior work by building models to predict traits related to litter quality and recalcitrance from reflectance spectra. Compared to previous studies, we calibrate and validate our models using more samples from more species. This breadth allows us to make predictions over a wide range of trait values and test how well models predict both intra- and interspecific trait variation. We also measured spectra from the same samples both while intact and ground. Existing models to predict leaf litter traits require the destructive and time-consuming step of grinding the tissue, which motivated us to test whether spectra of whole, intact litter could also yield accurate estimates. We predicted that ground-leaf litter spectra would provide more accurate estimates of chemical traits than intact-leaf litter spectra, as most comparisons on non-senesced leaves have shown (Serbin et al. 2014; Couture et al. 2016; Kothari et al. 2023b). On the other hand, we predicted that intact-leaf litter spectra would outperform ground-leaf litter spectra for estimating structural traits like LMA because grinding destroys the leaf structure (Kothari et al. 2023b).

Finally, we verified whether our litter chemistry estimates could capture known differences among species. We predicted that conifer litter would have greater LMA, recalcitrant carbon, and C_mass_ and lower N_mass_ than broadleaf litter. These patterns are thought to explain much of why conifer litter tends to show slower initial decomposition rates than broadleaf litter (Cornelissen et al. 1999; Cornwell et al. 2008; Prescott 2010).

## Methods

### Samples

During fall 2018, we collected senescent leaf litter from the Forests and Biodiversity experiment (FAB1) at Cedar Creek Ecosystem Science Reserve (East Bethel, MN, USA; Grossman et al. 2017). In May 2013, one-to two-year-old trees were planted in a 0.5 × 0.5 m grid within plots that varied in their species richness (1, 2, 5, or 12 species) and composition. The species pool included eight broadleaf angiosperms and four needleleaf conifers (Table 1). We left out one species (*Juniperus virginiana*) from sampling because its scaly needle morphology meant that individual trees seldom produced enough litter at once for spectroscopic or chemical analyses. To get enough samples of another species (*Acer negundo*), we also collected material from mature trees in a residential yard in Minneapolis, MN, USA 50.0 km from FAB1.

**Table 1:** Details of species in the Forests and Biodiversity (FAB1) experiment. All broadleaf species listed are deciduous angiosperms, and all needleleaf species are evergreen conifers. We did not sample *J. virginiana*. Twelve of the *A. negundo* samples come from a residential yard rather than FAB1. One *B. papyrifera* sample included in the tally below was omitted from ground litter analyses due to lack of tissue.

| Species | Code | Functional group | # samples |
| --- | --- | --- | --- |
| *Acer negundo* | ACNE | Broadleaf | 27 |
| *Acer rubrum* | ACRU | Broadleaf | 28 |
| *Betula papyrifera* | BEPA | Broadleaf | 36 |
| *Juniperus virginiana* | JUVI | Needleleaf | n/a |
| *Pinus banksiana* | PIBA | Needleleaf | 20 |
| *Pinus resinosa* | PIRE | Needleleaf | 31 |
| *Pinus strobus* | PIST | Needleleaf | 26 |
| *Quercus alba* | QUAL | Broadleaf | 37 |
| *Quercus ellipsoidalis* | QUEL | Broadleaf | 33 |
| *Quercus macrocarpa* | QUMA | Broadleaf | 34 |
| *Quercus rubra* | QURU | Broadleaf | 33 |
| *Tilia americana* | TIAM | Broadleaf | 17 |
| Total |  |  | 322 |

We collected litter samples as close as we feasibly could to the time of senescence using both litterfall traps on the ground and hand-collection of senesced leaves from trees. We only took leaves from trees when the petiole had a complete abscission layer and the leaf could be detached with a gentle pull (Chapin & Moilanen 1991). Because tree species dropped their leaves at different times, we collected litter repeatedly from September to November 2018 to ensure that the litter we collected had recently senesced. To avoid leaching we discarded litter samples from traps after rain, although nitrogen and carbon leaching tends to be low even under extreme wetting (Schreeg et al. 2013).

We collected 322 litter samples, seeking to represent the variation in leaf traits within each species (Table 1). While our sampling did not explicitly account for the species richness and composition treatments in FAB1, we collected leaves from each species across neighborhoods and crown positions to enhance intraspecific variation. Each sample included 1.5-3.0 g of tissue and could comprise one leaf or multiple similar leaves. When a single trap contained more than 2.5-3.0 g of leaves from a single species, we divided them into two or more samples by keeping together leaves that we judged by eye to be more visually similar. We dried each sample in a dark shed at 40 °C for at least three days, and then stored them in a cool, dark room until spectral and trait measurements.

### Spectral data collection

We measured reflectance spectra of each dried sample, both intact and ground, using a PSR+ 3500 full-range (350-2500 nm) spectrometer (Spectral Evolution, Lawrence, MA, USA). For intact-leaf spectra, we used the Spectral Evolution leaf clip assembly. We used a built-in 99% Spectralon white reflectance standard to calibrate readings before each sample. Among broadleaf species, many of the leaves curled in on themselves during senescence and drying; when possible, we sought to measure spectra in relatively flat portions of the leaf to minimize specular reflection (Petibon et al. 2021). Among needleleaf species, we made mats by laying out needles in a single layer; because of their shape, there often remained small gaps or areas of overlap between needles. We discarded spectra with large discrepancies in sensor overlap regions, or (particularly for needleleaf species) with very high (>70%) or very low (<30%) peak reflectance, which often indicated that the needle mat was highly non-uniform. There remained at least three reflectance spectra for each sample, and more for most of the broadleaf samples that had multiple leaves.

After removing their petioles, we ground leaf samples by placing them in individual plastic vials with ball bearings and shaking them with a paint shaker for several hours. To measure ground-leaf spectra, we used the Spectral Evolution benchtop reflectance probe and glass-windowed sample trays. Before measuring each sample, we measured a white Spectralon panel placed in a sample tray for calibration. For sample measurements, we put at least 0.6 g of leaf powder into a sample tray. Preliminary tests showed that adding more material did not change the spectra, suggesting that transmittance was close to zero. The benchtop probe pressed the loose powder into an even pellet. We measured three spectra per sample— turning the sample tray 120° between the first two measurements, and loosening and mixing the powder between the second and third measurements.

The spectrometer automatically interpolated the spectra to 1 nm resolution. We performed all subsequent processing using *spectrolab v. 0.0.18* (Meireles et al. 2023) in *R v. 3.6.1* (R Core Team, 2019). Because the 350-400 nm and 2400-2500 nm regions could be noisy, we trimmed each spectrum to 400-2400 nm. For intact leaves only, we corrected discontinuities at the overlap region between the Si and first InGaAs detectors (970-1000 nm) using the *match_sensors()* function, and spline-smoothed only this region using the *smooth_spline()* function. Finally, we averaged all intact spectra and all ground spectra for a given sample. As a metric of measurement consistency, we calculated the mean spectral angle between measurements of the same sample (Kruse et al. 1993). Intact measurements (median across samples: 6.8°; 2.5^th^-97.5^th^ percentile: 2.9°-20.4°) showed much less consistency than ground measurements (median: 0.6°; 2.5^th^-97.5^th^ percentile: 0.2°-3.2°).

### Leaf mass per area measurements

Researchers risk underestimating resorption efficiency when they fail to account for leaf mass loss during resorption. To avoid this risk, many studies either express nutrient content of non-senesced and senesced leaves on a per-area basis, or equivalently include a mass loss correction factor (van Heerwaarden et al. 2003). We measured LMA of each litter sample to enable conversion between a per-mass and per-area basis. We did not attempt to quantify or correct for the smaller bias sometimes caused by leaf shrinkage (van Heerwaarden et al. 2003).

For broadleaf species, we could not measure the area of whole leaves because many were curled and brittle. Instead, we used a hole punch to take four to six 0.3 cm^2^ disks from each dried leaf sample. We sought to capture the variation within the sample, including veins. We calculated LMA as the total mass divided by the total area of all disks. This method has generally been shown to provide accurate estimates of the LMA of whole leaves (Perez et al. 2020).

For needleleaf species, we measured the mass and area of five needles per sample. We estimated area as the length times the maximum width of the needle measured using digital calipers. We then measured the mass of each needle and calculated LMA as the total mass divided by the total area.

### Chemical trait measurements

We measured leaf carbon and nitrogen concentration (C_mass_ and N_mass_) on ground, oven-dried samples using dry combustion gas chromatography performed by an elemental analyzer (Costech ECS 4010 Analyzer, Valencia, California, USA). We converted C_mass_ and N_mass_ to an area basis (C_area_ and N_area_) by multiplying them with LMA.

We measured carbon fractions on ground, oven-dried samples using an ANKOM 200 Fiber Analyzer (Ankom Technology, Macedon, New York, USA). The first digestion in a hot neutral detergent solution washes out plant solubles. The second digestion in a hot acidic detergent solution further washes out bound proteins and hemicellulose (henceforth ‘hemicellulose’ for simplicity). The remaining fraction comprises cellulose and acid-unhydrolyzable residue (AUR) such as lignin (henceforth ‘recalcitrant carbon’). We estimated each component using the changes in sample mass between steps. Because the paint shaker ground our samples into a very fine powder, we had to use small-pored ANKOM F58 fiber bags, which cannot be used in the final acid detergent lignin digestion to measure AUR concentration alone. We also did not determine the ash concentration of the samples.

We removed some trait values prior to statistical analyses because of clear issues during trait data collection—for example, the rupture of sample bags while measuring carbon fractions. Unlike some other studies (e.g., Gillon et al. 1999), we only removed samples due to known issues with trait measurements, not simply because they had large prediction residuals during statistical analyses.

### Statistical analyses

We modeled the relationship between traits and spectra using partial least-squares regression (PLSR), which is well-suited to handle datasets that include many collinear predictors (Burnett et al. 2021). PLSR reduces the full matrix of spectra to a smaller number of orthogonal latent components that best explain the variation in the variables to be predicted. We implemented our PLSR models in *R* package *pls v. 2.7.1* (Mevik et al. 2019).

Previous studies using a PLSR modeling framework have often restricted the range of wavelengths used to predict certain traits in order to select bands that, based on prior research or known absorption features, may be most causally linked to the traits in question (e.g., Serbin et al. 2014). In our main set of analyses, we used the full spectrum (400-2400 nm) to predict each trait on the grounds that biochemical changes during senescence may alter the relationship between integrative functional traits like N_mass_ and specific optical features. However, we also explored two ways of restricting the spectral range for intact-leaf spectra only. First, we built models using only 400-1000 nm (‘VIS/NIR models’) given that many less-expensive spectrometers only measure this range. Second, we built models using only 1300-2400 nm (‘SWIR models’). The pigment composition of leaves changes rapidly during senescence, including breakdown products (‘brown pigments’) which may absorb up to 1300 nm (Fourty et al. 1996). The SWIR models could perform better in cases where pigments are poor indicators of the overall composition of the leaf.

Researchers may transform reflectance of ground-leaf spectra to pseudoabsorbance (*-log R*; Blackburn 1998; Serbin et al. 2014) or its derivatives, which may somewhat linearize the relationship between traits and spectral features and make those features more prominent. We found in preliminary tests that these transformations did not substantially improve model performance. Likewise, brightness normalization of intact-leaf spectra (Feilhauer et al. 2010) did not have a strong influence on model performance.

We divided our samples into calibration (75%) and validation (25%) datasets, stratified such that the proportions of each species were identical in each dataset. We fit models for each trait on the complete calibration dataset, using 10-fold cross-validation to select the optimal number of model components to use in further analyses while avoiding overfitting. We selected the smallest number of components whose cross-validation root mean squared error of prediction (RMSEP) was within one standard deviation of the very lowest RMSEP at any number of components. We used these models to calculate the variable importance in projection metric (VIP) for each trait (Wold et al. 1994) but we do not report their accuracy.

We developed our main set of models using a resampling procedure in which we divided the calibration data at random 200 times into 70% training and 30% testing subsets (Burnett et al. 2021). During each resampling iteration, we trained a model for each trait on the 70% and assessed its performance (RMSE and *R^2^*) on the remaining 30%, using the previously determined number of components for that trait. This procedure left us with an ensemble of 200 models for each trait, which could be used both to make trait predictions and to examine the variability of model performance under random variation in training data.

We used these ensembles of models to predict traits from the validation (25%) dataset. We quantified model performance for each trait by calculating *R^2^* and root mean squared error (RMSE) between the predictions (averaged for each validation sample across the 200 estimates) and the measured values. We also report the %RMSE, calculated as the RMSE divided by the 2.5% trimmed range of the measured values in the validation data (Kothari et al. 2023a).

## Results

There was a high level of variation in each trait both across the entire dataset and within species. For example, N_mass_ varied nearly 11-fold; in contrast, C_mass_ only varied about 1.5-fold (Table 2). In spite of the high level of intraspecific variation, especially among broadleaf species, species identity alone could account for between 46.0% (N_area_) and 93.7% (LMA) of the variation in each trait. In general, needleleaf species had much higher LMA, C_mass_, recalcitrant carbon, and lower N_mass_ than broadleaf species (Table S1), consistent with trends in non-senescent leaves (Serbin et al. 2014; Kothari et al. 2023a). Consequently, we see correlations among pairs of leaf economic traits across the whole dataset (for log(LMA) and log(N_mass_), *R^2^* = 0.592; *p* < 10^−15^), but a weaker correlation among just needleleaf samples (*R^2^* = 0.255; *p* < 10^−5^) and none among just broadleaf samples (*R^2^*= 0.011; *p* = 0.111).

**Table 2:**
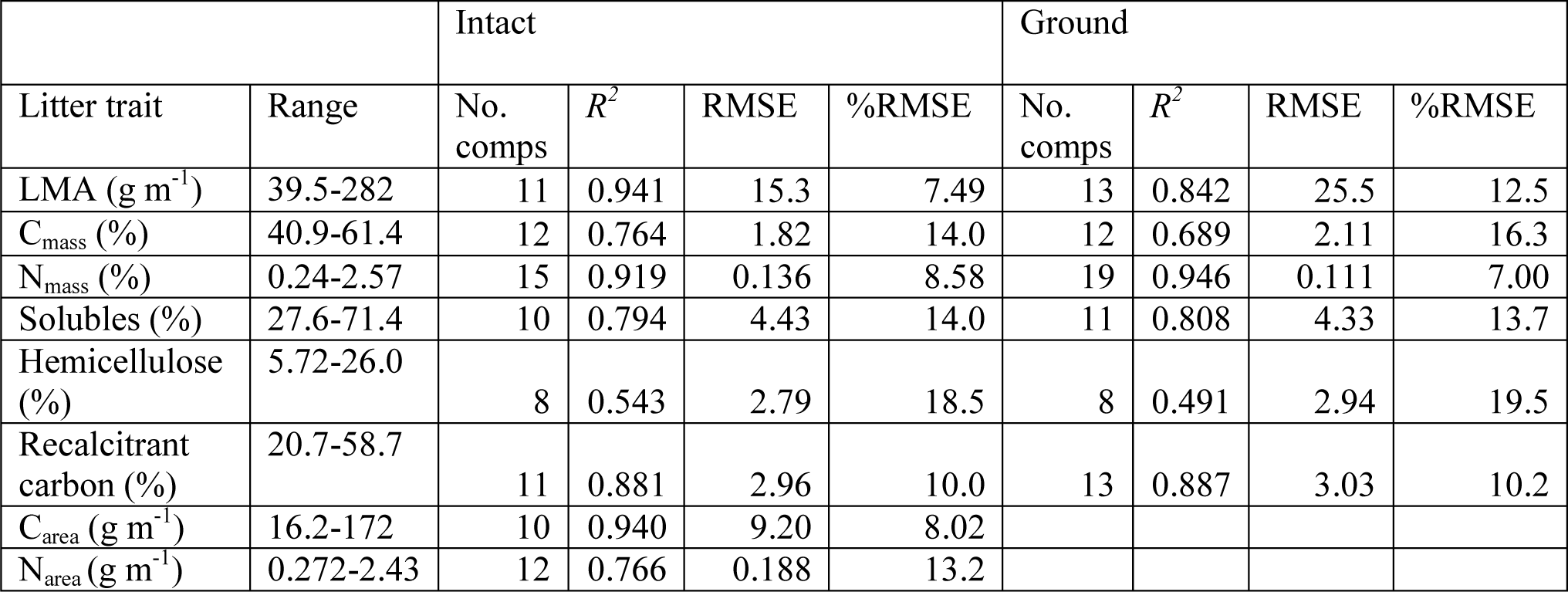
Summary statistics of PLSR model validation based on full (400-2400 nm) intact- and ground-leaf spectra. The range column shows the complete trait range across the full dataset. %RMSE is calculated as RMSE divided by the 2.5% trimmed range of data within the validation data set.

Among both ground-leaf and intact-leaf spectra, the coefficient of variation (CV) was highest in the visible range and decreased into the near-infrared range (NIR; Fig. 1). For intact-leaf spectra only, the CV once again increased into the short-wave infrared range (SWIR). There was a global maximum CV around 440-490 nm for intact spectra, shifted to lower wavelengths for ground spectra; there were also local maxima centered at around 670-675 nm for both kinds of spectra. Both maxima lie near absorption peaks of chlorophylls, and reflectance at 670 nm is particularly sensitive to variation in chlorophyll at low levels, such as during late senescence (Gitelson & Merzlyak 1996).

**Fig. 1:**
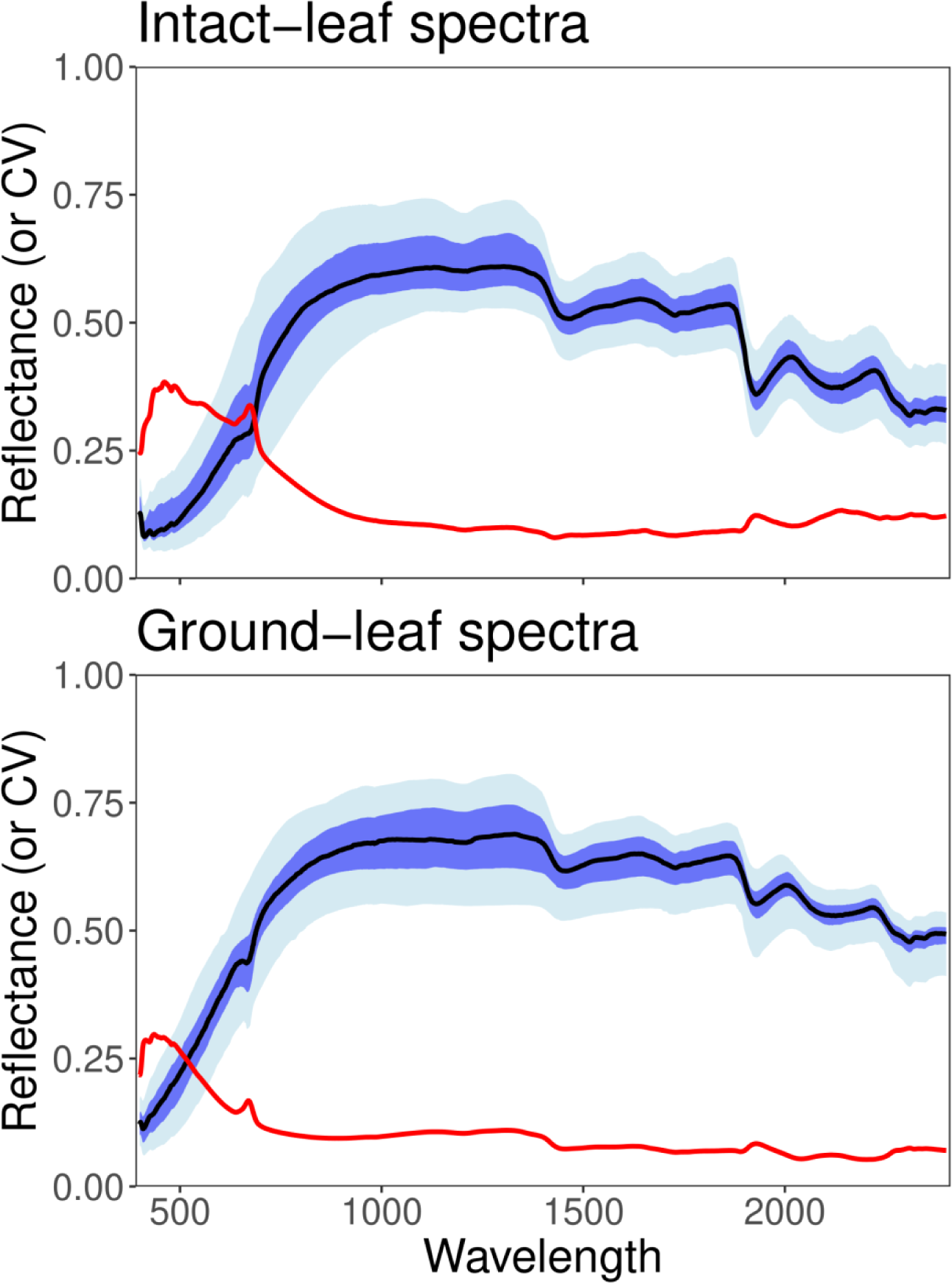
Distributions of spectral reflectance and its coefficient of variation (CV) among intact (top) and ground (bottom) leaf litter samples. The black line represents the median reflectance at each wavelength, flanked by the 25-75 percentile region (dark blue), and the 2.5-97.5 percentile region (light blue). The solid red line shows the coefficient of variation among spectra at each wavelength.

Each of our traits could be predicted with moderate-to-high success from full intact-leaf and ground-leaf spectra (Figs. 2-4; Table 2). The models for N_mass_ and (especially for intact-leaf spectra) LMA achieved the highest calibration and validation accuracy, followed by recalcitrant carbon, solubles, C_mass_, and finally hemicellulose (Table 2). There was no general tendency for estimation of chemical traits to be better using intact- or ground-leaf spectra, but estimation of LMA was better using intact-leaf spectra. Because we used LMA to convert from a mass to an area basis, we built models predicting N_area_ and C_area_ (nitrogen and carbon on a per-area basis) only from intact-leaf spectra (Fig. 4). The C_area_ model (%RMSE = 8.02) had better performance than the C_mass_ model (%RMSE = 14.0), but the N_area_ model (%RMSE = 13.2) had worse performance than the N_mass_ model (%RMSE = 8.58; Table 2). Jackknife analyses showed that traits for which the ensemble model performance was poorest—such as hemicellulose, C_mass_, and N_area_—also had the greatest variability in model performance (Figs. S1 and S2).

**Fig. 2:**
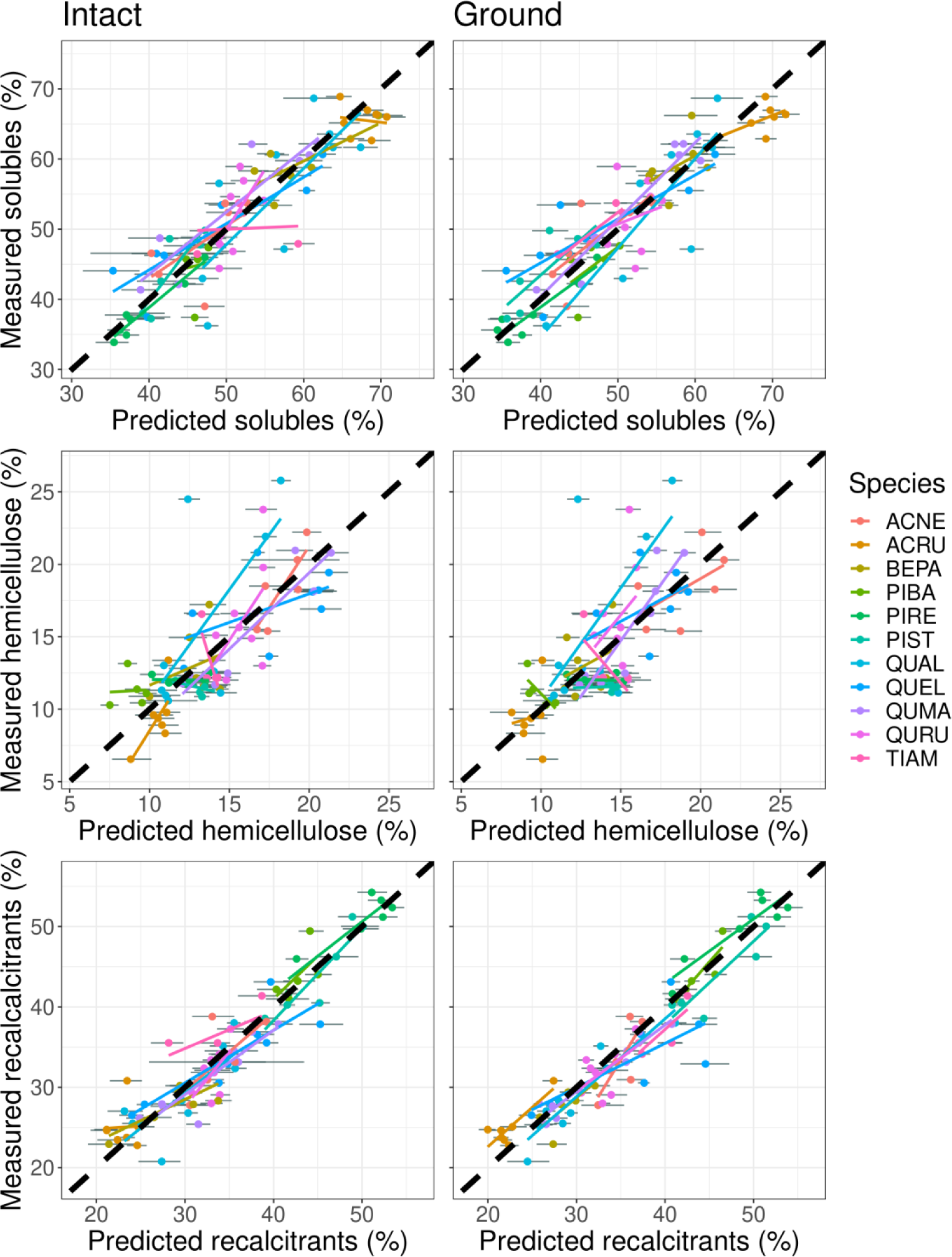
Validation results for predictions of solubles, hemicellulose, and recalcitrant carbon from intact-leaf litter spectra (left) and ground-leaf litter spectra (right). In each panel, a separate OLS regression line is shown for each of the 11 species, overlaid on top of the thick dashed 1:1 line. The error bars for each data point are 95% intervals calculated from the distribution of predictions based on the model coefficients from the 200 jackknife iterations. See Table 1 for species codes.

**Fig. 3:**
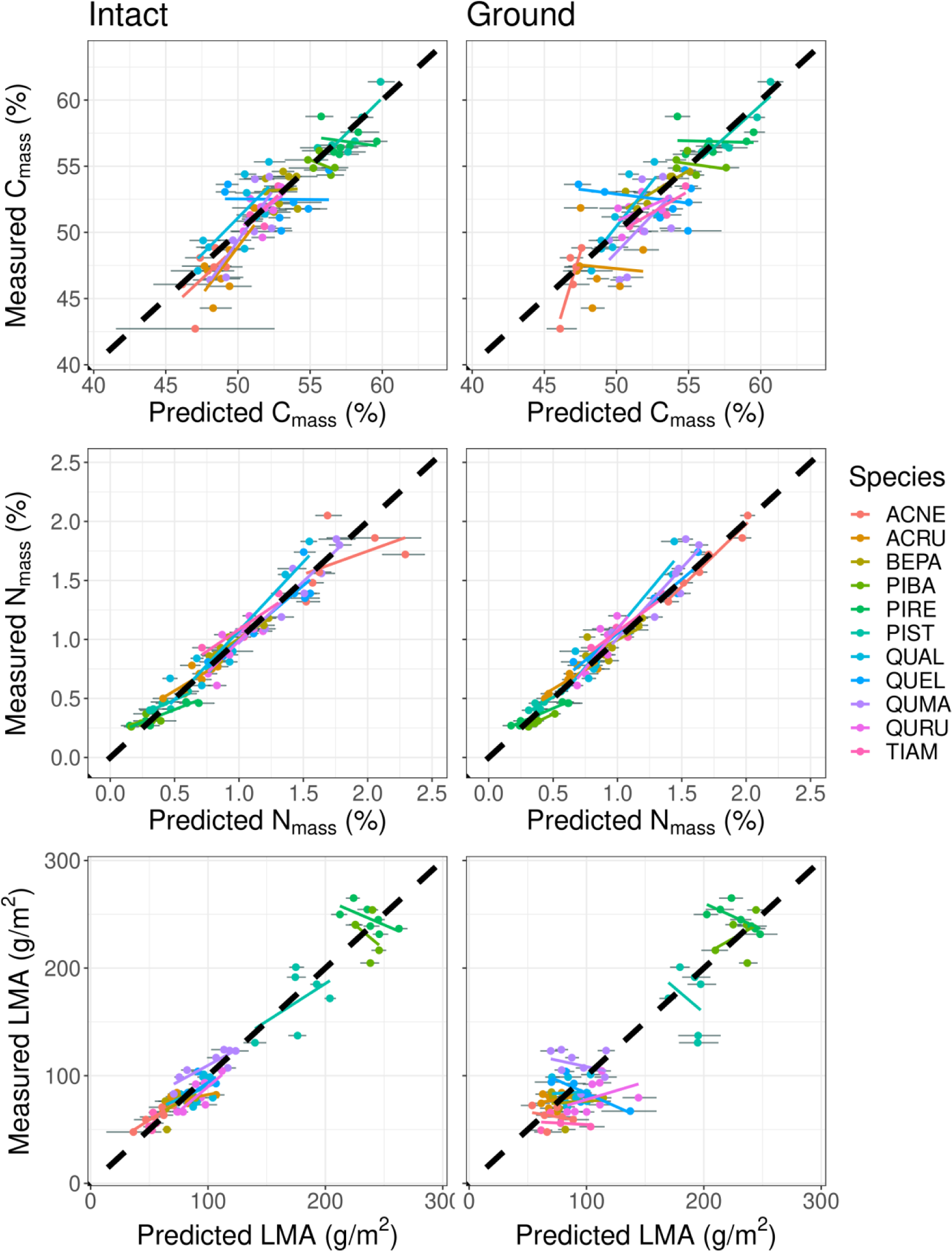
Validation results for predictions of C_mass_, N_mass_, and LMA from intact-leaf litter spectra (left) and ground-leaf litter spectra (right). Regression lines and error bars are as in Fig. 2. See Table 1 for species codes.

**Fig. 4:**
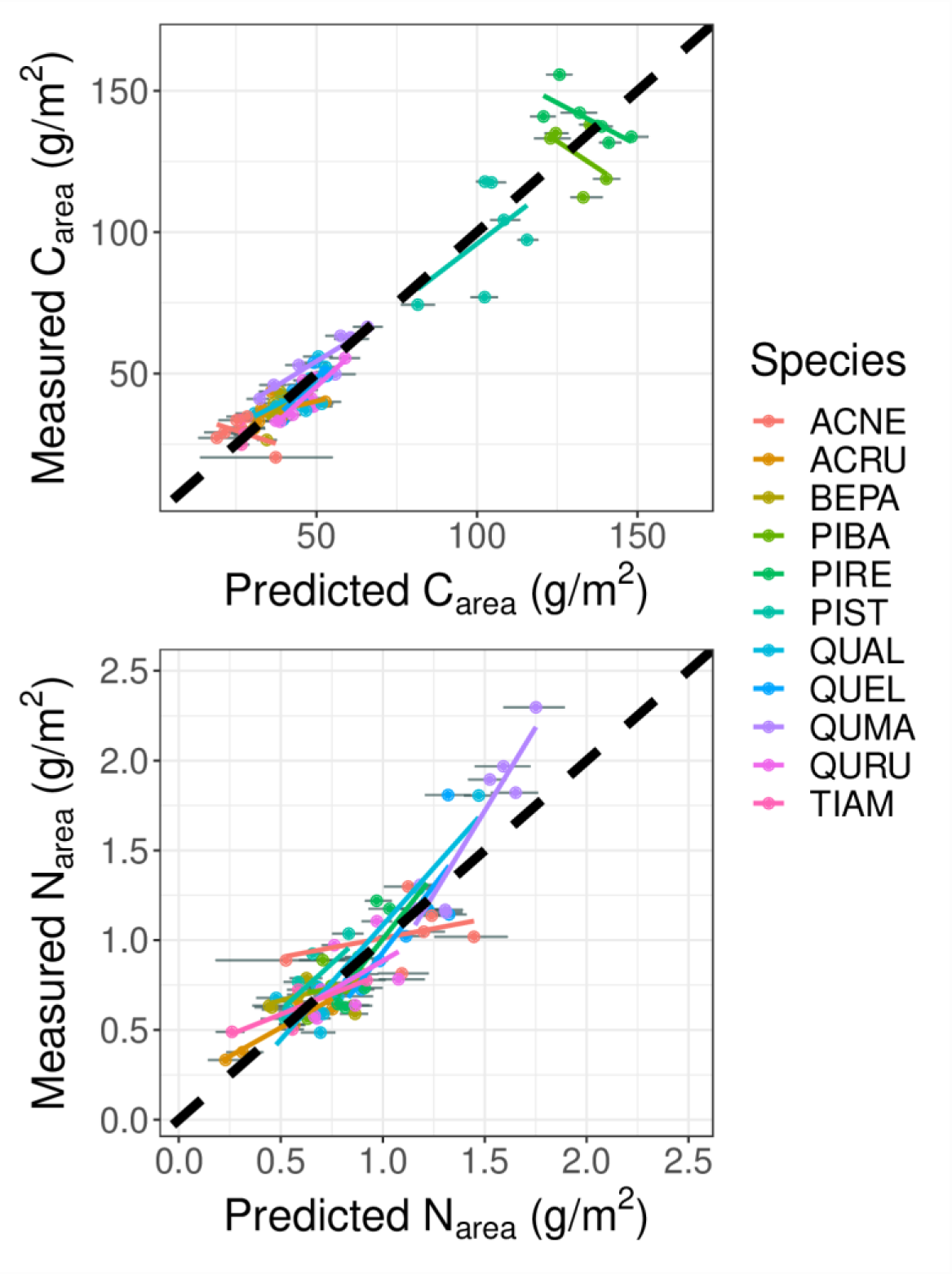
Validation results for predictions of C_area_ and N_area_ from intact-leaf litter spectra. Regression lines and error bars are as in Fig. 2. See Table 1 for species codes.

For every trait prediction model trained on the full spectrum, the VIP metric also showed a global maximum around 677 nm (intact) or 672 nm (ground; Fig. 5), next to one of the aforementioned chlorophyll absorption peaks. This pattern suggests that the amount of chlorophyll remaining at abscission may be closely related to many traits, especially N_mass_. More generally, the visible range was most important for predicting all traits, and the NIR range was also important for LMA. There were further peaks in the SWIR around 1430-1450 nm (for intact leaves), 1910-1950 nm and 2130-2220 nm. In fresh tissue, variation in reflectance around 1910-1950 nm is often interpreted as a measure of absorption by water, but the trace amounts of water remaining in our dried samples likely have no direct biological relevance. In general, reflectance in the visible range and parts of the SWIR range seemed most informative in predicting nutrients and carbon fractions in litter.

**Fig. 5:**
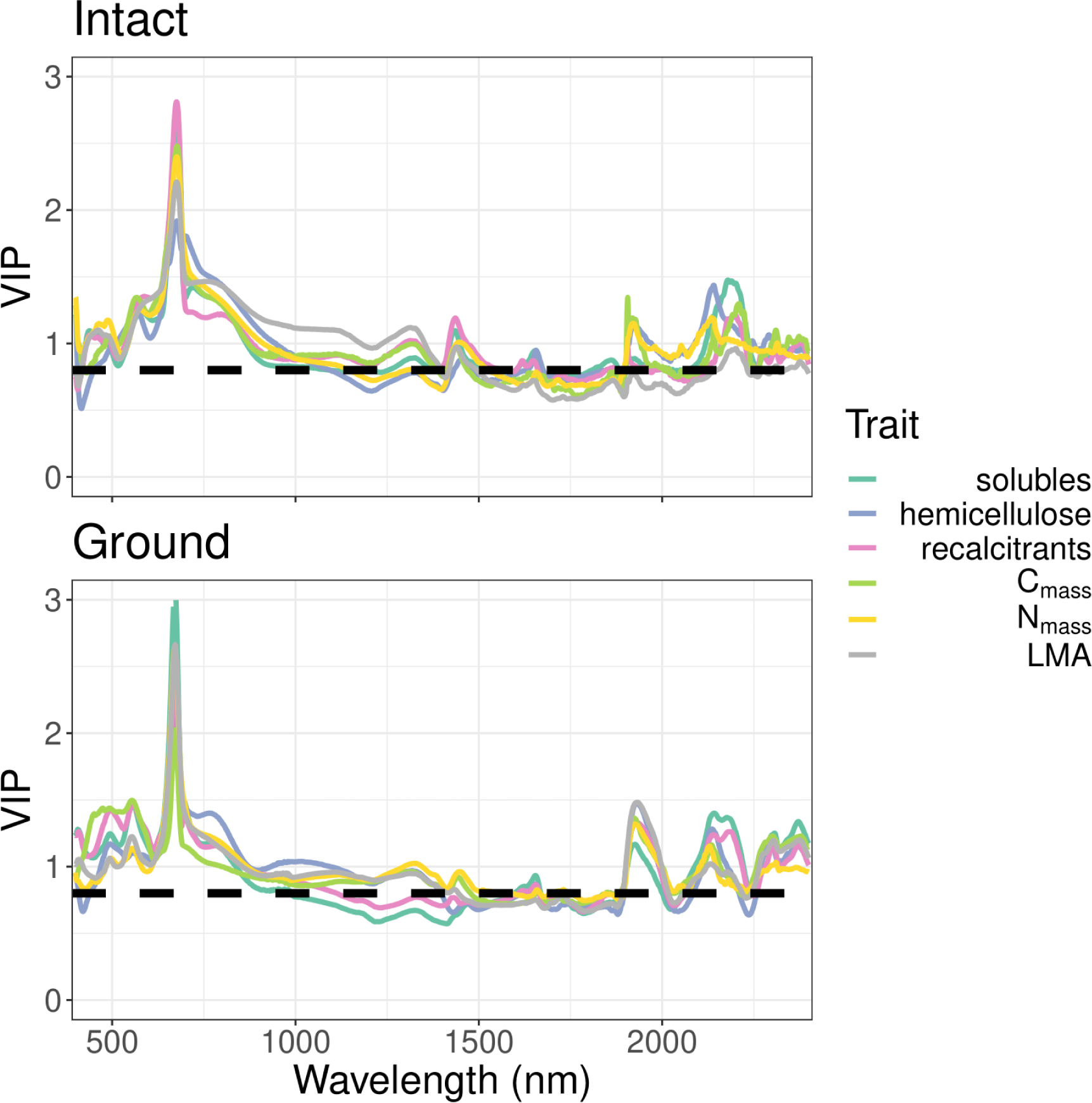
The variable importance of prediction (VIP) metric in intact (top) and ground (bottom) models. The dashed black line represents a threshold of 0.8 suggested as a rough heuristic of importance by Burnett et al. (2021).

Our VIS/NIR models for intact leaves had considerably worse performance than our full-range models for every trait, particularly N_mass_, N_area_, and solubles. By contrast, our SWIR models had very similar performance to the full-range models (Table S2). Both sets of restricted-range models had a similar rank-ordering of model performance across traits to each other and the full models. Moreover, the VIP metric for these models revealed that the regions that were most important for these predictions were mainly the same bands in their respective regions that were important for the full-range models; however, for the VIS/NIR models, very low wavelengths close to 400 nm also had high importance.

## Discussion

We show that PLSR models trained on reflectance spectra can accurately predict several leaf litter traits related to nutrient cycling across temperate broadleaf and needleleaf tree species. For each trait, a single model captures the variation among and within our 11 species. The models’ predictions capture well-known ecological differences among species—for example, that needleleaf species have more recalcitrant carbon and lower N_mass_ than broadleaf species. Importantly, intact-leaf models are about as accurate as ground-leaf models for most traits, which suggests that multiple traits can be estimated from a single measurement that takes only seconds.

### Model performance and interpretation

The hierarchy of which traits can be predicted best mirrors results from non-senesced leaves, including fresh ones (Asner et al. 2011; Nunes et al. 2017; Kothari et al. 2023a). Leaf mass per area can be predicted very well from intact-leaf spectra in particular, which has been shown in both fresh leaves (Serbin et al. 2019) and in pressed, dried leaves (Costa et al. 2018; Kothari et al. 2023b). Predictions of LMA from ground leaves are worse, but still quite good (Serbin et al. 2014). N_mass_ can also be predicted nearly perfectly from intact- and ground-leaf spectra (Richardson & Reeves 2005; Serbin et al. 2014). Recalcitrant carbon, solubles, and C_mass_ all can be predicted with *R^2^* > 0.65, while the smallest carbon fraction (hemicellulose) shows the worst performance (Kothari et al. 2023b). For chemical traits, intact- and ground-leaf models had only small and idiosyncratic differences in performance.

Our comparisons of variation among replicate spectral measurements within samples imply that grinding homogenizes the variation in the leaf and allows more consistent measurements. This consistency may owe to several factors. Veins and other features cause intact leaves to vary in their chemical makeup across the laminar area, which may be hard to capture representatively. Likewise, if chemical traits change with depth through the laminar cross-section, the optical features of deeper layers may be obscured in part by shallower layers. The geometries of uneven needle mats and curled-up leaves may also reduce consistency by increasing variation in anisotropic surface reflectance (Petibon et al. 2021). Nevertheless, there was no strong, consistent indication that ground-leaf spectra are necessarily better than intact-leaf spectra for predicting chemical traits. This result deviates from the few previous comparisons of intact- and ground-leaf spectra (Serbin et al. 2014; Couture et al. 2016; Kothari et al. 2023b). It is unclear why the theoretical considerations in favor of ground-leaf spectra do not amount to much in practice here; we speculate that information about structure might, depending on the context, either confound or indirectly aid the prediction of chemical traits.

In contrast to chemical traits, LMA was clearly better predicted from intact-leaf spectra. Given that grinding destroys the leaf structure, it may be perplexing that ground-leaf spectra predict structural traits like LMA at all (Serbin et al. 2014; Kothari et al. 2023b). The relationship between ground-leaf spectra and LMA is likely indirect, mediated through correlations between LMA and chemical traits that retain clear optical signatures in ground tissue (Nunes et al. 2017; Kothari & Schweiger 2022). Since trait correlations within species often differ from those among species (Anderegg et al. 2018), such effects can explain why a model trained across many species may fail to capture intraspecific variation (Kothari & Schweiger 2022). Indeed, in our dataset, ground-leaf spectra do a poor job of predicting intraspecific variation in LMA; species-specific slopes of measured vs. predicted values are often far from 1.

Changing the basis of nitrogen and carbon from mass to area also changes the accuracy of estimates. While N_mass_ is predicted more accurately than N_area_, C_mass_ is predicted less accurately than C_area_. Across samples, N_mass_ correlates negatively and C_mass_ correlates positively with LMA, driven in large part by the split between broadleaf (low LMA) and needleleaf (high LMA) species. As a result, multiplying by LMA to convert N and C to an area basis flattens interspecific variation in N, but enhances it in C. For example, the percentage of variation explained by species identity alone increases for carbon (*R^2^* from 0.721 for C_mass_ to 0.945 for C_area_), but decreases for nitrogen (*R^2^* from 0.786 for N_mass_ to 0.460 N_area_). Absorption features that differentiate species or functional groups can aid estimates of C_area_ much more than N_area_, whether or not they are causally related to those traits. Area- and mass-based traits often have different uses: there is a strong justification for using area-based nutrients for studying resorption (van Heerwaarden et al. 2003), while mass-based traits are more often used for explaining decomposition (Cornwell et al. 2008), and a ‘normalization-independent’ basis has been proposed for making sense of trait covariance (Osnas et al. 2018). In general, the basis of estimation should be determined by these factors rather than the apparent accuracy of the models.

Visible-range wavelengths where chlorophyll and other pigments absorb had high importance in our main set of full-range models. However, removing these wavelengths (‘SWIR models’) did not decrease performance while removing the SWIR range (‘VIS/NIR models’) did. It seems that pigments are neither essential for predicting litter chemistry, nor do they confound it. Although the bands where they absorb are assigned high importance, much of the information may be redundant with that of the SWIR range, where many leaf macromolecules absorb. However, those SWIR features are not fully redundant with the information about pigments and leaf structure captured by the VIS/NIR range. The spectrum of leaf or litter samples could remain more stable above 1300 nm over the long term (Kothari et al. 2023b), so the SWIR models may also transfer better when sample preparation and storage practices are inconsistent.

### Advancing spectroscopic estimation of litter traits

Compared to conventional measurements, spectroscopic estimates of plant tissue functional traits are easy and fast to generate and can be non-destructive. Spectrometers are costly, but measuring each spectrum is (on the margin) nearly free. Once samples have been collected and organized, measuring each sample’s intact-leaf spectrum takes under a minute. In contrast, both ground-leaf spectra and conventional measurements require grinding leaves, which is time-consuming and destructive. Using our protocol, we could measure the spectrum of a single ground sample (including sample tray preparation and cleanup) in 5-7 minutes, not including grinding time. In contrast, we found that doing both elemental and carbon fraction analysis takes about 25 minutes of operator time per sample. The time saved through spectroscopic trait estimation grows as the number of traits of interest increases.

Many studies that leverage spectroscopy to address ecological questions use models tailored to the particular species and tissues under study (e.g., McTiernan et al. 2003; Hobbie 2005; Fortunel et al. 2009). While this practice allows researchers more confidence in their trait estimates, it requires them to spend time and money doing the manual trait measurements needed for calibration, which they must repeat for any new study system they adopt. Spectroscopic trait estimation is unlikely to serve as a complete substitute for standard measurements unless we can create models that are demonstrably accurate across a wide range of species and conditions.

The good validation performance of our models does not guarantee accurate trait estimates from any set of litter samples. To take one example, consider the aforementioned pattern that the ground-leaf model for LMA often performs poorly within species, despite its strong performance across species. If it were used to study intraspecific variation in litter traits, the model’s estimates could be useless or even misleading. Moreover, when models simply ‘learn’ to exploit any features that happen to correspond with LMA among species within a given dataset—without respect to their causal relationship with LMA—they may also transfer poorly to new species (Kothari & Schweiger 2022). This case is closely linked to the more general problem in machine learning of distinguishing spurious and invariant correlations (Arjovsky et al. 2019). As the volume of available spectral data increases, taking care to assess out-of-sample generalization and borrowing tools from machine learning could help us train better spectral models and make more realistic judgments about their performance (Kattenborn et al. 2022; Cherif et al. 2023).

Recent studies have attempted to tackle the challenge of building global trait-spectra models. For example, Serbin et al. (2019) show that a single model can accurately predict the LMA of fresh, non-senesced leaves across 11 sites from the tropics to the Arctic. It remains to be seen whether any single model could predict traits like N_mass_ or LMA from leaf tissue across many species at all stages of senescence and decomposition. The predictive success of global models may rely on the constrained patterns of trait covariation among non-senesced leaves—for example, along continua from sun to shade leaves or conservative to acquisitive leaves (Osnas et al. 2018)—which may help in estimating traits that are not themselves associated with strong absorption features (Nunes et al. 2017; Kothari & Schweiger 2022). However, the dramatic shifts in leaf chemistry and cellular structure during senescence (Keskitalo et al. 2005)—including pigment degradation and nutrient resorption—could weaken some of these relationships among traits, or between traits and their optical features. If so, it could be hard to build a global model for senesced leaves that achieves the predictive success of global models for non-senesced leaves.

Some evidence suggests that litter traits do in fact predictably covary with each other, and with non-senesced leaf traits (Freschet et al. 2010; Freschet et al. 2012; Jackson et al. 2013). But while most fresh green leaves look alike, each species’ leaves may senesce in its own way. For example, some of our species (*A. rubrum*, *Q. alba*, *Q. ellipsoidalis*, *Q. rubra*) tend to produce reddish anthocyanins during senescence, but the rest do not. The breakdown of certain metabolites during senescence also creates a complex mixture of brown pigments (Fourty et al. 1996), which are poorly described and may vary from species to species. Such variation could make the relationship between traits and optical features more contingent for senescent foliage.

Even setting aside these concerns, model generality may also be complicated by differences in sample preparation (e.g., grind size) and instruments, which may affect the spectrum in subtle but important ways (Foley et al. 1998; Petibon et al. 2021). Overcoming these challenges may require investing in protocol development, or in trying algorithms that are more flexible than PLSR in ‘learning’ the complex forms of biological variation in heterogeneous datasets. In the meantime, it would be prudent for researchers who plan to use existing spectral models in new systems to take some conventional trait measurements in order to validate model performance.

Finally, we note that plants produce and shed many kinds of tissue besides leaves, and these other litter sources also contribute to nutrient cycling through resorption and decomposition (Lü et al. 2012). Reflectance spectroscopy may help measure nutrient-related traits in other tissues. For example, fine root decomposition is an important but little-understood part of nutrient cycling; fine roots may contribute nearly half of annual litter inputs in forests (Freschet et al. 2013) and vary considerably in resorption efficiency (Freschet et al. 2010). Elle et al. (2019) showed that PLSR models built using near-infrared reflectance spectroscopy can predict fine root lignin, which increases recalcitrance (See et al. 2019; but see, in contrast, Sun et al. 2018). Continuing to develop models for other plant tissues could make it easier to study how plants alter nutrient cycling in a holistic way. Likewise, nutrients other than nitrogen often limit or co-limit growth and decomposition, and tools to estimate them quickly and cheaply could advance the study of plant nutrient economies. However, estimates of some micronutrients from reflectance spectra may be unreliable (Nunes et al. 2017; Kothari et al. 2023a), and other methods may prove more useful (e.g., X-ray fluorescence spectroscopy; van der Ent et al. 2018).

Ultimately, whether accurate global models for predicting litter traits from spectra can be built is an empirical question that can only be settled by amassing a wide variety of data. We take an initial step towards this long-term goal by showing that reflectance spectroscopy can provide accurate litter trait estimates across many temperate tree species that vary widely in their litter traits. This line of research shows great promise in relieving some of the major limitations toward understanding plant functional ecology in an ecosystem context.

## Acknowledgements

The University of Minnesota, including Cedar Creek, is located on the traditional and contemporary Dakota and Ojibwe land. Many members of the Cavender-Bares Lab and the UMN Physiological Ecology Discussion Group contributed to discussions that improved this paper. We give particular thanks to Laura Williams, Artur Stefanski, Sam Reed, and Habacuc Flores-Moreno for useful discussions and help in data collection, as well as Rebecca Montgomery for contributions to the FAB1 experiment. The research was supported by an Alexander & Lydia Anderson Grant from University of Minnesota and by the National Science Foundation’s funding of Cedar Creek LTER under DEB #1234162. Spectral measurements were conducted as part of an NSF/NASA Dimensions of Biodiversity project (DEB #1342778) and the NSF ASCEND Biology Integration Institute (DBI #2021898). SK was supported by an NSF Graduate Research Fellowship (Grant No. 00039202) and a UMN Doctoral Dissertation Fellowship.

## Data availability

All spectral and trait data will be shared on the EcoSIS repository upon acceptance for publication. Analysis code will be archived on Zenodo. (A current version of the analysis code is available at: https://github.com/ShanKothari/senesced-trait-models.) Models will be uploaded to the Data Repository of U of M (DRUM).

## Author contributions

SK and JCB conceived the project with input from SEH. SK collected the spectral and trait data and conducted all data analysis. SK wrote the first draft of the manuscript, and all authors contributed to further revisions.

## Supplementary Material

**Table S1:**
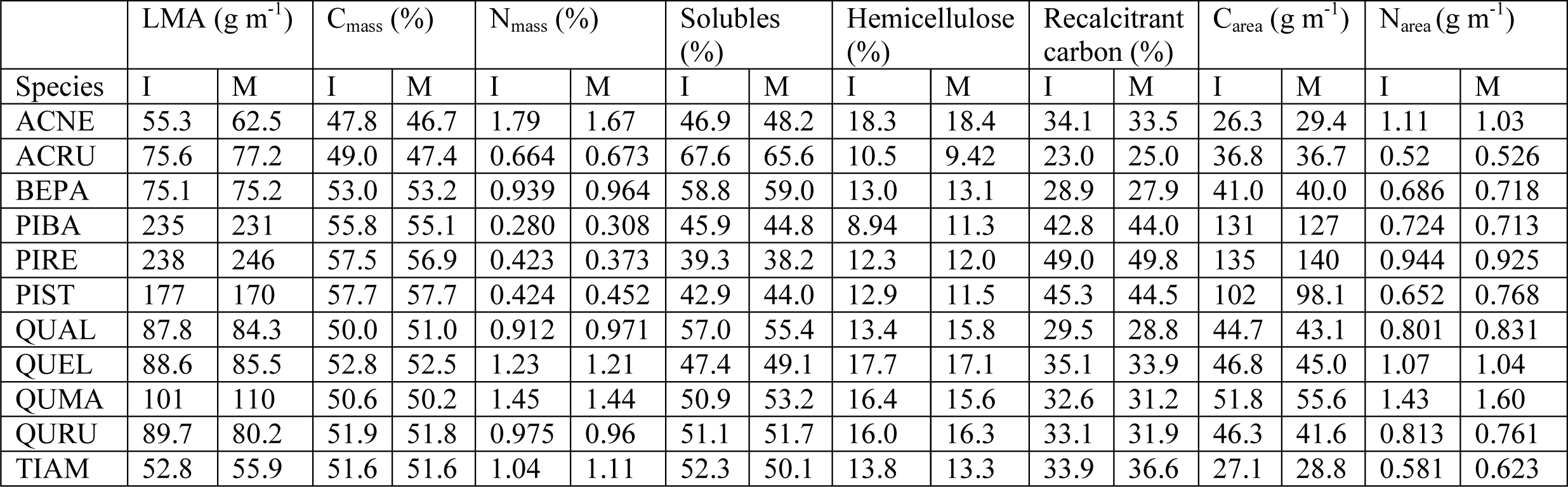
Species means of predicted values from intact-leaf models (I) and measured values (M) within the validation dataset for each of the eight leaf litter traits under consideration. See Table 1 in main text for species codes.

**Table S2:** Summary statistics of PLSR model validation based on intact- and ground-leaf spectra restricted to the VIS/NIR range (400-1000 nm) or to part of the SWIR range (1300-2400 nm). %RMSE is calculated as RMSE divided by the 2.5% trimmed range of data within the validation data set.

|  | VIS/NIR |  |  |  | SWIR |  |  |  |
| --- | --- | --- | --- | --- | --- | --- | --- | --- |
| Trait | No.<br>comps | $R^2$ | RMSE | %RMSE | No.<br>comps | $R^2$ | RMSE | %RMSE |
| LMA ($\text{g m}^{-1}$) | 10 | 0.871 | 23.1 | 11.3 | 7 | 0.939 | 15.6 | 7.63 |
| C<sub>mass</sub> (%) | 10 | 0.663 | 2.15 | 16.6 | 6 | 0.647 | 2.21 | 17.0 |
| N<sub>mass</sub> (%) | 12 | 0.712 | 0.25 | 15.8 | 11 | 0.939 | 0.12 | 7.57 |
| Solubles (%) | 8 | 0.613 | 5.92 | 18.7 | 6 | 0.782 | 4.58 | 14.4 |
| Hemicellulose (%) | 8 | 0.371 | 3.30 | 21.9 | 6 | 0.555 | 2.76 | 18.3 |
| Recalcitrant carbon (%) | 12 | 0.789 | 3.91 | 13.2 | 9 | 0.869 | 3.07 | 10.4 |
| C<sub>area</sub> ($\text{g m}^{-1}$) | 10 | 0.873 | 13.6 | 11.9 | 7 | 0.942 | 9.12 | 7.95 |
| N<sub>area</sub> ($\text{g m}^{-1}$) | 9 | 0.588 | 0.249 | 17.4 | 10 | 0.782 | 0.186 | 13.0 |

**Fig. S1:**
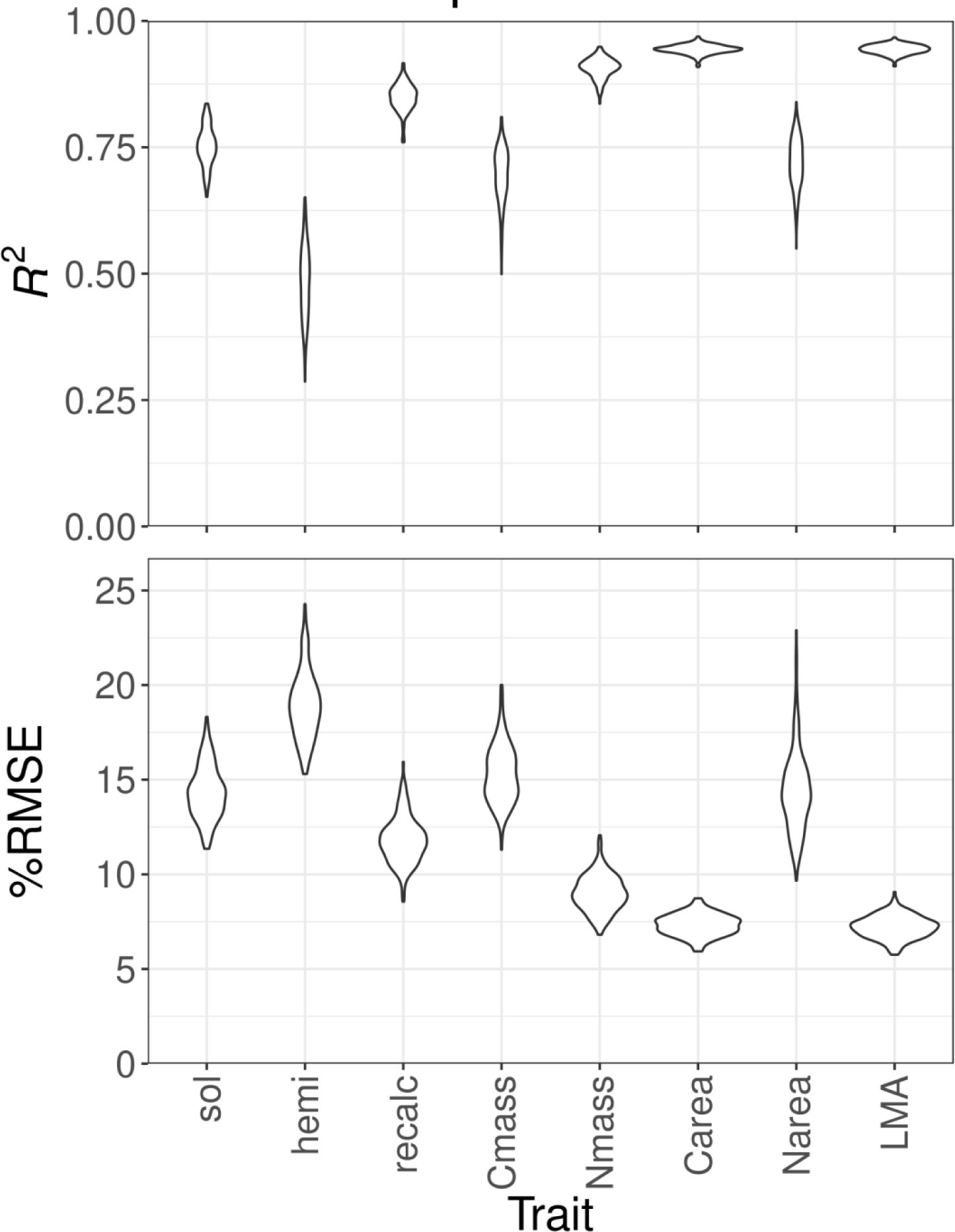
Intact-leaf litter spectral model performance statistics for each trait based on 200 jackknife iterations from the calibration data set. %RMSE is calculated as RMSE divided by the 2.5% trimmed range of data within the testing data set. In this case, the testing data set for each iteration is the 30% of data randomly chosen from within the calibration set not used for training the model in that iteration. Abbreviations: sol = solubles, hemi = hemicellulose, recalc = recalcitrant carbon.

**Fig. S2:**
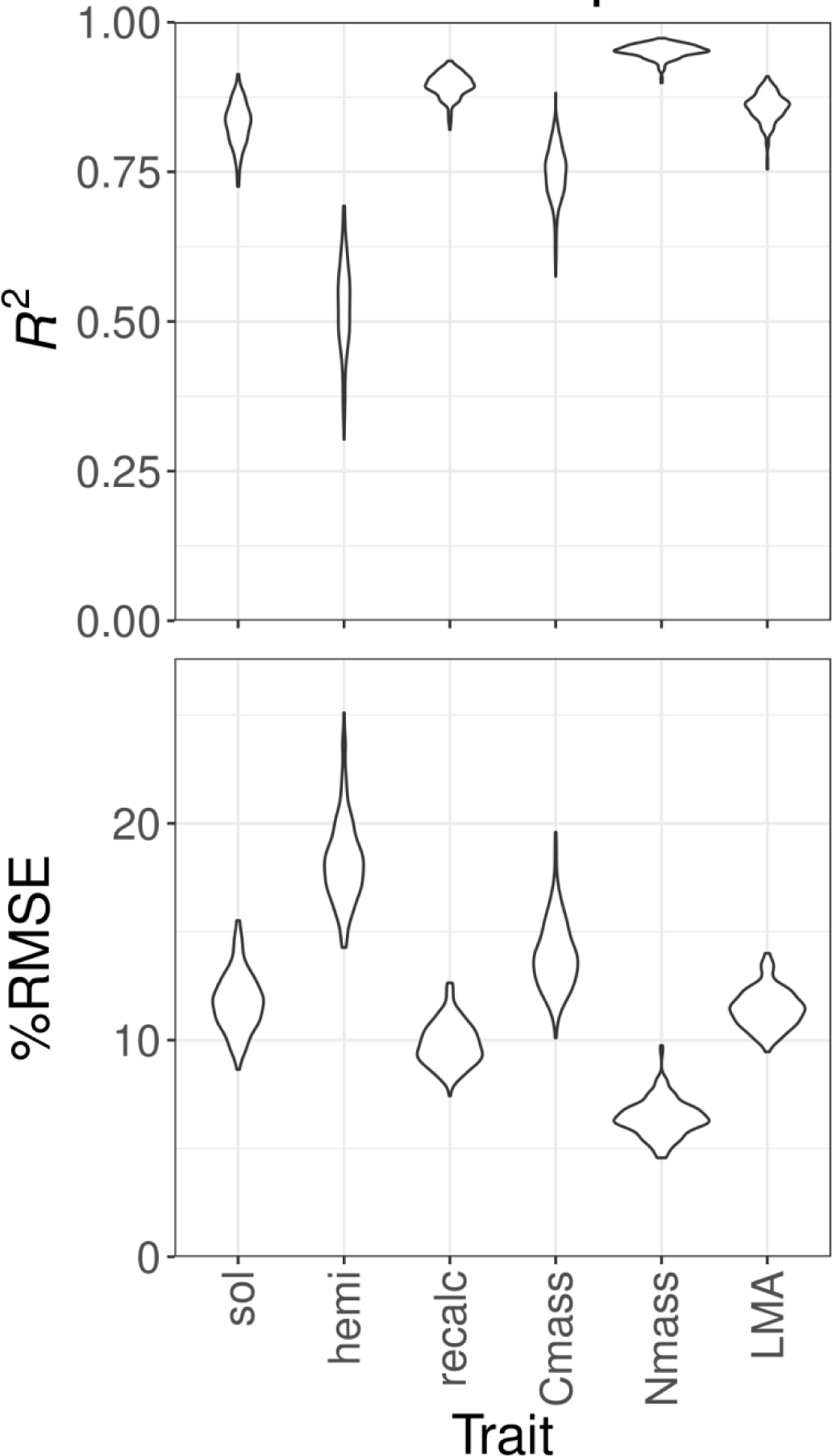
Ground-leaf litter spectral model performance statistics for each trait based on 200 jackknife iterations from the calibration data set. %RMSE is calculated as RMSE divided by the 2.5% trimmed range of data within the testing data set. In this case, the testing data set for each iteration is the 30% of data randomly chosen from within the calibration set not used for training the model in that iteration. Abbreviations: sol = solubles, hemi = hemicellulose, recalc = recalcitrant carbon.

**Fig. S3:**
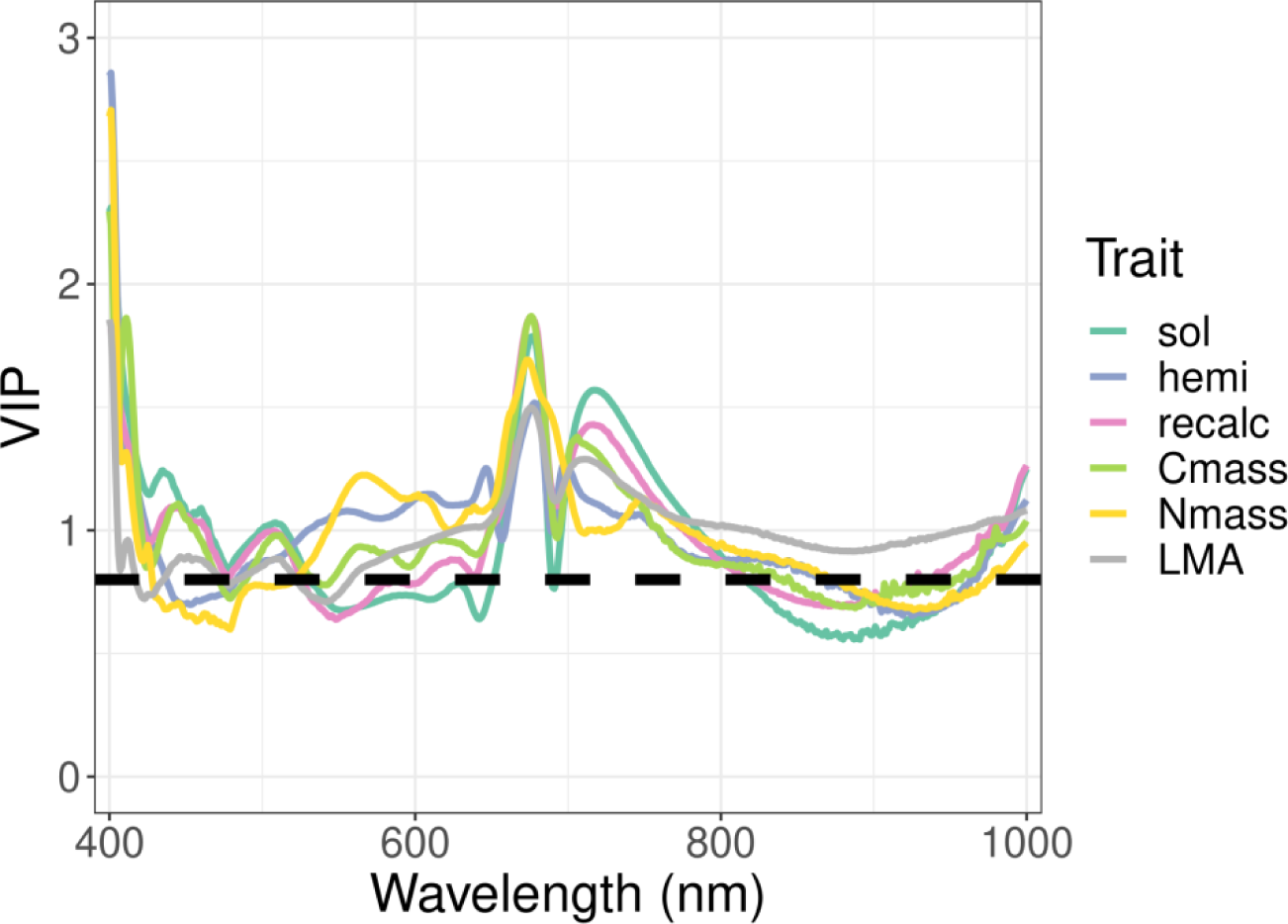
The variable importance of prediction (VIP) metric in intact-leaf VIS/NIR models. The dashed black line represents a threshold of 0.8 suggested as a rough heuristic of importance by Burnett et al. (2021). Abbreviations: sol = solubles, hemi = hemicellulose, recalc = recalcitrant carbon.

**Fig. S4:**
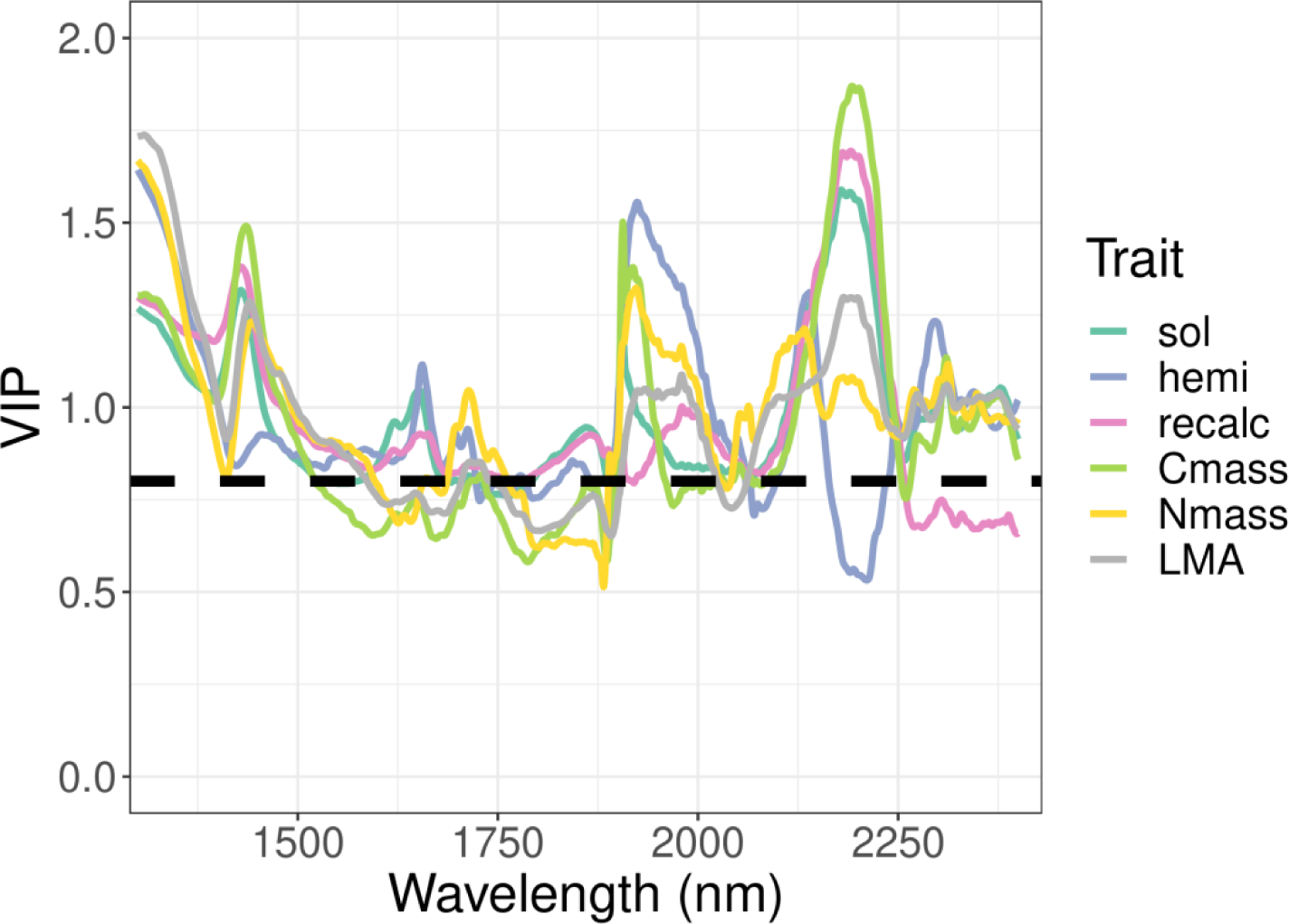
The variable importance of prediction (VIP) metric in intact-leaf SWIR models. The dashed black line represents a threshold of 0.8 suggested as a rough heuristic of importance by Burnett et al. (2021). Abbreviations: sol = solubles, hemi = hemicellulose, recalc = recalcitrant carbon.

